# Barcoded SARS-CoV-2 viruses define the impact of time and route of transmission on the transmission bottleneck in a Syrian hamster model

**DOI:** 10.1101/2024.06.08.597602

**Authors:** Reed Trende, Tamarand L. Darling, Tianyu Gan, David Wang, Adrianus C.M. Boon

**Author notes:** Corresponding author. Adrianus C. M. Boon.

## Abstract

The transmission bottleneck, defined as the number of viruses that transmit from one host to infect another, is an important determinant of the rate of virus evolution and the level of immunity required to protect against virus transmission. Despite its importance, SARS-CoV-2’s transmission bottleneck remains poorly characterized, in part due to a lack of quantitative measurement tools. To address this, we adapted a SARS-CoV-2 reverse genetics system to generate a pool of >200 isogenic SARS-CoV-2 viruses harboring specific 6-nucleotide barcodes inserted in ORF10, a non-translated ORF. We directly inoculated donor Syrian hamsters intranasally with this barcoded virus pool and exposed a paired naïve contact hamster to each donor. Following exposure, the nasal turbinates, trachea, and lungs were collected, viral titers were measured, and the number of barcodes in each tissue were enumerated to quantify the transmission bottleneck. The duration and route (airborne, direct contact, and fomite) of exposure were varied to assess their impact on the transmission bottleneck. In airborne-exposed hamsters, the transmission bottleneck increased with longer exposure durations. We found that direct contact exposure produced the largest transmission bottleneck (average 27 BCs), followed by airborne exposure (average 16 BCs) then fomite exposure (average 8 BCs). Interestingly, we detected unique BCs in both the upper and lower respiratory tract of contact animals from all routes of exposure, suggesting that SARS-CoV-2 can directly infect hamster lungs. Altogether, these findings highlight the utility of barcoded viruses as tools to rigorously study virus transmission. In the future, barcoded SARS-CoV-2 will strengthen studies of immune factors that influence virus transmission.

## INTRODUCTION

A defining feature of the SARS-CoV-2 pandemic has been the continual emergence of variants despite the low mutation rate of SARS-CoV-2 compared to other ssRNA viruses (1.3 × 10^-6^ mutations per nucleotide per replication cycle^1, 2^, less than 10% the mutation rate of influenza A, HCV, or HIV^3^). One underexplored factor that may partially explain this seeming contradiction is the transmission bottleneck, defined as the number of viruses that spread from one host to infect another. The transmission bottleneck can impact the rate of viral evolution via several mechanisms. First, wider transmission bottlenecks may help a virus overcome pre-existing immunity as the host will have to neutralize more viruses to prevent infection^4^. Second, wider transmission bottlenecks increase the prevalence of mixed-strain infections and probability of recombination across strains^5, 6^. Third, wider transmission bottlenecks increase the efficiency of variant selection by increasing the amount of viral diversity transferred between hosts^6, 7^. Conversely, narrow transmission bottlenecks only transfer a small amount of genetic diversity between hosts, increasing the likelihood that fit variants fail to transmit due to purely stochastic factors. Narrow bottlenecks can also fix deleterious mutations through a process known as Muller’s ratchet, though in viral infections this is likely to only impact isolated transmission chains rather than global viral evolution^7–9^. Supplementing these theoretical arguments that wider bottlenecks aid viral evolution, empirical evidence has shown that wider transmission bottlenecks helped preserve antiviral-resistant influenza strains in a ferret transmission model^10^. Altogether, these arguments highlight that better understanding the transmission bottleneck has important implications for SARS-CoV-2 evolution.

Several studies have measured the transmission bottleneck of SARS-CoV-2 in humans by tracking shared genetic variants between known transmission pairs. Similar experimental strategies has been previously deployed with success for mutation-prone viruses such as influenza that produce many intrahost genetic variants, suggesting a narrow average bottleneck (1-5 virions)^11–13^. However, estimated bottlenecks for individual transmission pairs can be extremely variable; McCrone et al^11^ estimated that 95% of transmission bottlenecks are ≤3 virions, but identified one pair with an estimated bottleneck >200 virions. When applied to SARS-CoV-2, iSNV studies also consistently measure a narrow transmission bottleneck (1-5 virions) between people^13–17^. However, the low frequency of SARS-CoV-2 intra host variants makes these studies very sensitive to the parameters used in iSNV calling^15^, and prevents the identification of transmission pairs based solely on sequence identity^16, 18^. Further, these studies rest on several assumptions that are often untested, such as that detected iSNVs do not impact viral fitness and that there are no lower respiratory tract-specific iSNVs, as this site is often not sampled in humans. Additionally, these studies have insufficient power and precision to measure how variables like duration of exposure and host immune status impact the SARS-CoV-2 transmission bottleneck. These shortcomings necessitate the development of animal models where the transmission bottleneck can be studied quantitatively.

Barcoded (BC) viruses containing a fitness-neutral genetic barcode have been effectively used to study virus transmission for an array of respiratory and non-respiratory viruses^19^, including influenza^20–25^, HIV and SIV^26–32^, Zika virus^33–36^, West Nile Virus^37^, and Coxsackie virus^38^. BC viruses overcome many of the shortcomings of measuring transmission by using iSNVs. Genetic diversity increases the power of both BC virus pools and iSNVs to detect transmission events, but while iSNV diversity is limited by intrinsic viral properties, BC virus pools can be made arbitrarily genetically diverse, enabling much more precise measurement of the transmission bottleneck. Additionally, the fitness-neutrality of the BC design can be validated. Finally, the use of a predefined pool of BCs can greatly increase the sensitivity and specificity of detecting BCs and thus transmission events.

Given the need to better understand SARS-CoV-2 transmission and the utility of BC viruses, we designed, generated, and validated a pool of >200 BC SARS-CoV-2 viruses. To study SARS-CoV-2 transmission, donor hamsters were inoculated with the pool of BC SARS-CoV-2, contact hamsters were exposed to the donors, and the number of BCs transmitted was quantified. We then evaluated the impact of duration and route of exposure, demonstrating that longer exposure times and more direct routes of exposure widen the SARS-CoV-2 transmission bottleneck. Interestingly, we also identified tissue-specific BCs in the nasal turbinate, trachea, and lungs, suggesting that transmitted SARS-CoV-2 can directly seed each tissue. Altogether, these studies generated a pool of BC SARS-CoV-2, a novel and powerful tool for the study of SARS-CoV-2 transmission and leveraged this tool to study SARS-CoV-2 spread within and between hosts in unprecedented detail.

## RESULTS

### Generation and Validation of Genetically Barcoded SARS-CoV-2 Viruses

The SARS-CoV-2 transmission bottleneck in humans and animal models remains incompletely characterized. To define the transmission bottleneck, we generated genetically barcoded SARS-CoV-2 viruses, allowing us to quantify the number of unique infection events in the recipient host. Using an established SARS-CoV-2 reverse genetics system^39^, we inserted a 9 nucleotide sequence harboring a 6 nucleotide barcode (BC) (GNNGNNGNN) in ORF10, a non-translated ORF near the 3’ UTR in the prototypic SARS-CoV-2 virus. We also introduced a D614G mutation in the Spike gene of SARS-CoV-2. This mutation has been associated with enhanced transmission in humans and various animal models^40^. Individual BC viruses were isolated, expanded, and titered (see Methods for details) (**Fig. 1A**). To verify the fitness-neutrality of our BC insert *in vitro*, we compared the growth kinetics of a representative panel of 11 individual BC viruses to a sequence-confirmed WT-D614G stock. All tested BC viruses had similar growth kinetics to the stock (**Fig. 1B**), indicating that the BC insert does not attenuate the virus *in vitro*. Based on these results, we pooled >200 BC viruses and WT-D614G based on equal infectious viral titers (hereafter referred to as the “BC Pool”).

**Figure 1:**
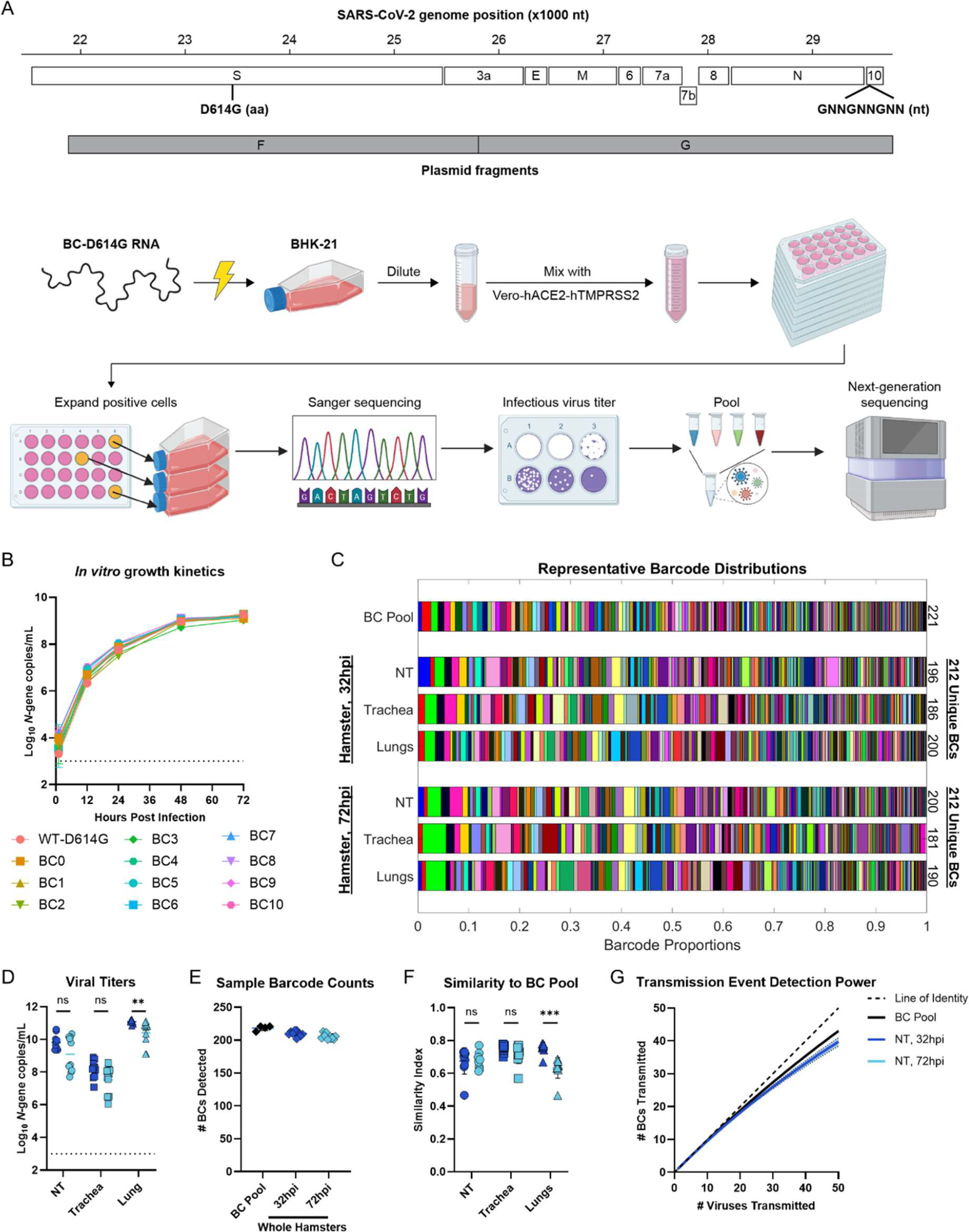
Generation and validation of genetically barcoded SARS-CoV-2 viruses. **(A)** Schematic of barcoded SARS-CoV-2 virus generation. We utilized an established reverse- genetics system comprised of seven DNA segments spanning the prototypic SARS-CoV-2 genome with the spike D614G mutation (segment F) and a 9nt insert containing a 6nt BC within ORF10 (segment G). The *in vitro*-transcribed barcoded viral genomes were electroporated into BHK-21 cells, mixed 1:1 with Vero-hACE2-hTMPRSS2 cells, and plated into multiple 24-well plates. Virus-positive wells were individually expanded on Vero-hTMPRSS2 cells, Sanger sequenced, titered by plaque assay, and pooled and the resulting distribution was analyzed by next-generation sequencing (NGS). (**B**) *In vitro* growth characteristics of barcoded viruses. Vero- hACE2-hTMPRSS2 cells were inoculated with an MOI of 0.01 and supernatant was collected 12, 24, 48 and 72 hours later and used to quantify virus titer by RT-qPCR. Each color represents a unique BC virus. (**C-H**) Naïve male hamsters were inoculated with 10^5^ PFU of the BC virus pool and nasal turbinate, trachea, and lung tissues were collected 32 or 72 h later (n=10 per group). The barcode sequence and frequency was determined by NGS. (**C**) A representative image depicting the frequency and number of BC detected in the BC SARS-CoV-2 pool and in each tissue of inoculated hamsters collected at 32 or 72 hpi. Each color represents a unique BC and bar width is proportional to the BC’s relative frequency. The total number of unique BCs detected in each tissue and in the entire animal is shown on the right side of the graph. (**D**) Virus titers in the nasal turbinate, trachea and lungs of BC SARS-CoV-2 inoculated hamsters 32 (dark blue) and 72 hpi (light blue), measured by RT-qPCR. (**E**) Number of unique BCs detected in at least one tissue of each infected hamster. (**F**) Similarity index between the BC pool and different respiratory tissues of the inoculated hamsters 32 and 72 hours post inoculation. (**G**) Average number of unique BCs transmitted for a given number of virus transmission events. Results are the average following 10,000 transmission events for each number of viruses. NT = nasal turbinate. Dotted lines indicated limit of detection. The line is the geometric mean (**D**) or average (**E-F**) of the data. Data was log-transformed where necessary and analyzed by an unpaired t-test followed by a Holm-Šídák test (**** *P* < 0.0001, *** *P* < 0.001, ** *P* < 0.01, * *P* < 0.05, ns = not significant). The results are from 2 independently repeated experiments with five hamsters each.

To measure the BC distribution in the BC pool, viral RNA was extracted, and the BC region was amplified by a cycle-limited RT-PCR. The resulting amplicons were sequenced by next-generation sequencing, and the barcode sequences present and their associated frequency was determined (see Methods for details). This entire pipeline was independently repeated 4 times, yielding distributions of 217-224 unique barcodes above a cutoff threshold (0.05%) (**Fig. 1C** and **S1A**), the vast majority of which (>94% of all barcodes) were detected in all runs (**Fig. S1B**). Note that, across all figures showing BC distributions in this paper, the same color represents the same BC, and all BCs are shown in the same left-to-right order. Further, BCs were detected at highly reproducible frequencies, as shown by the strong correlation in BC frequency across sequencing runs (**Fig. S1C**). Additionally, a similarity index^41–43^ that accounts for both the presence and relative frequency of BCs showed a very high degree of similarity between all sequencing runs (**Fig. S1D**). Based on these results, we defined a final list of BCs in our pool consisting of all BCs present in a majority of sequencing runs. Greater than 99% of barcode pairs were separated by a Hamming Distance >1 (**Fig. S1E**), minimizing the risk that sequencing errors could lead to false positive BC detections. Additionally, several measures of BC distribution quality were computed. Average richness (the number of BCs detected in a sample)^21^ was 218.5 (**Fig. S2A**); average evenness (a measure of BC distribution uniformity, range 0-1)^44^ was 0.96 (**Fig. S2B**); and average Shannon diversity (a metric that accounts for the number and uniformity of BCs)^45^ was 5.19 (**Fig. S2C**).

To assess BC virus fitness *in vivo*, we intranasally (IN) inoculated male hamsters with 10^5^ PFU of the BC pool of BC and collected nasal turbinates, trachea, and lung, 32 and 72 hrs post inoculation (hpi). RNA was extracted and virus titers were measured by RT-qPCR^46^ (**Fig. 1D**). The geometric mean viral RNA levels in the nasal turbinate, trachea, and lungs at 32 hpi were 6.6*10^9^, 1.4*10^8^, and 1.2*10^11^ *N*-gene copies/mL, respectively. At 72 hpi, nasal turbinate and trachea viral titers remained similar while lung titers dropped to 2.5*10^10^ *N*-gene copies/mL. All these titers are in line with previously reported hamster infections^40, 47^. Next, BC distributions in each tissue were measured by NGS as described above and examined. Qualitatively, BC distributions in all donor tissues are qualitatively similar to the BC Pool (**Fig. 1C**). Quantitatively, infected hamsters harbored 201-215 unique BCs in at least one respiratory tissue (**Fig. 1E**). All individual respiratory tissues also had rich, even, and diverse BC distributions (**Fig. S2A-C**). Additionally, all tissues had a high degree of similarity to the BC pool at 32 hpi (∼0.7, approximately as similar as two tissues from different donors) (**Fig. 1F**). Nasal turbinate and trachea similarity indices remained consistent to 72hpi, while in lungs the similarity indices dropped slightly (to 0.62, *P* <0.01) (**Fig. 1F**). We also calculated the geometric mean fold change in frequency for each barcode between the BC pool and each tissue at 32 and 72 hpi (**Fig. S2D**). Only 2 BCs expanded by >3-fold at both 32 and 72 hpi compared to the inoculum, and both only did so in a single tissue, suggesting that no BC virus has a meaningful competitive fitness advantage over the other viruses in the pool. This is further supported by comparing the fold change in BC frequency in hamsters between 32 and 72 hpi. Only 3 BCs expanded >3-fold *in vivo*, each did so in only a single tissue, and BC fold-change was not correlated across tissues (**Fig. S2E**).

To precisely measure the transmission bottleneck, BC virus pools should be sufficiently diverse so that the likelihood a given BC is transmitted more than once is low. To directly estimate the impact of barcode collisions on our transmission event detection power, we calculated the average number of unique BCs that would be transmitted for a given number of virus transmission events for our BC pool and for all inoculated hamster nasal turbinates (**Fig. 1G**). In the BC pool and hamster nasal turbinates, BC counts accurately measure transmission events when <30 viruses are transmitted and underestimate the transmission bottleneck by 5 or more when ≥30 viruses are transmitted. Collectively, these results demonstrate that our BC pool is sufficiently rich and diverse to measure a wide transmission bottleneck and that it retains its richness and diversity *in vivo*.

### Defining the Transmission Bottleneck in Airborne-Exposed Hamsters

Having functionally validated the pool of BC SARS-CoV-2 viruses in direct hamster infection, we proceeded to use it to quantitatively study SARS-CoV-2 transmission in an established airborne exposure model^48^. Briefly, male donor hamsters were inoculated IN with 10^5^ PFU of the BC pool. Twenty-four hours later, a naïve male contact hamster was exposed to the infected donor hamster for 8 hrs in a fresh cage. The donor and contact hamsters were physically separated by at least 2 cm by porous stainless steel, allowing air to flow from the infected donor to the contact hamster. Nasal turbinates, trachea, and lungs were collected from contact hamsters 72 hours post exposure (hpe) (**Fig. 2A**). RNA was extracted from each tissue and used to measure virus titers by RT-qPCR and BC distributions by NGS. Viral titers were consistently high across all collected contact tissues (**Fig. 2B**), demonstrating that the BC viruses can transmit via the airborne route in hamsters. Analysis of the BC sequence distributions revealed that in the five airborne-exposed contact hamsters, we detected an average of 23 unique BC per animal (range 13-42) (**Fig. 2C**). An average of 16 BCs were found in the lung (range 12-28), 14 in the trachea (range 5-28), and 12 in the nasal turbinates (range 4-20). While most BCs were shared between multiple respiratory tissues, tissue-specific BCs were identified in all respiratory tissues of most animals, suggesting unique infection events in both upper and lower airways of airborne-exposed hamsters (**Fig. 2C-D**).

**Figure 2:**
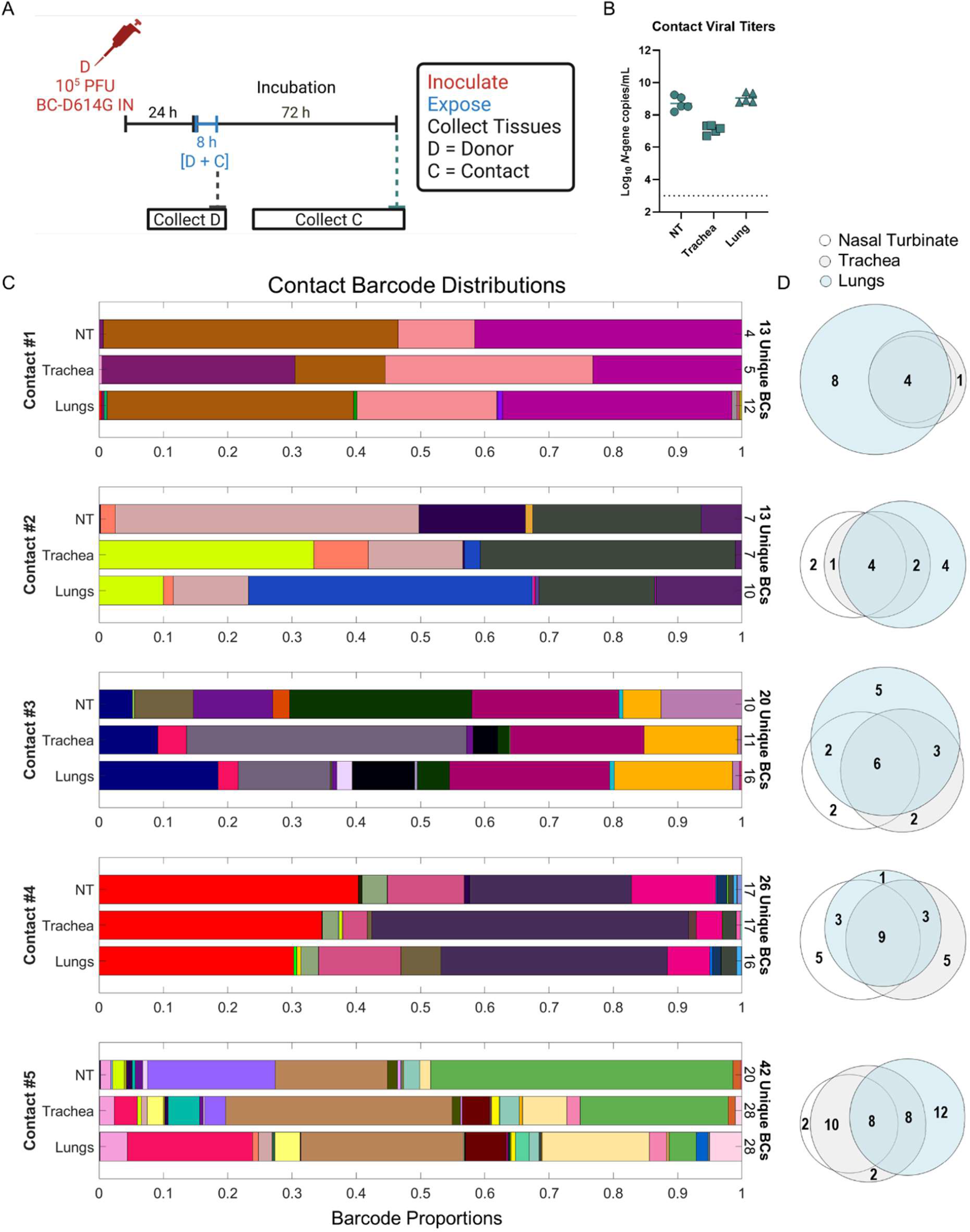
Defining the SARS-CoV-2 airborne transmission bottleneck. **(A)** Schematic of experimental design. Contact hamsters (n=5) were exposed for 8 hours to donor hamsters 24hrs after inoculation of 10^5^ PFU of the BC virus pool in our airborne exposure setup. Seventy-two hours later, nasal turbinate, trachea, and lung tissues were collected, and RNA extracted. (**B**) Virus titers, measured by RT-qPCR, in the nasal turbinate (circles), trachea (squares), and lung (triangles) of airborne exposed contact hamsters. (**C**) Barcode distributions in each tissue of all five contact hamsters. Each color represents a unique BC and bar width is proportional to the BC’s relative frequency. The total number of unique BCs detected in each tissue and in the entire animal is shown on the right side of the graph. (**D**) Proportional Venn diagram showing barcodes unique to and shared between the three respiratory tissues in each animal. Dotted lines indicated limit of detection. Bars indicate geometric mean. The results are from a single experiment with five donor and contact hamsters.

Virus transmission can be conceptualized as a series of bottlenecks, including but not limited to transfer to a new host and expansion within that host. A recent study by Holmes et al^22^ using BC influenza reported much higher BC counts in contact animals 1-2 days post-exposure than at later timepoints, demonstrating that expansion is a major bottleneck in influenza transmission. To assess whether expansion also imposes a bottleneck on SARS-CoV-2 transmission, we collected the respiratory tissues from airborne-exposed contact hamsters 24 and 48hpe (**Fig. S3A**). Shorter incubation periods in the contact hamsters resulted in more variable viral titers, though peak titers in each organ were generally similar (**Fig. S3B-C**). Importantly, unlike influenza, we did not observe higher BC counts at earlier timepoints (**Fig. S3D-E**). In fact, there was a trend towards lower BC counts at earlier timepoints. This strongly suggests that expansion is not a meaningful bottleneck in SARS-CoV-2 transmission. Interestingly, tissue-specific BCs were more prevalent at 24hpe compared to tissues collected 48 or 72hpe (**Fig. S3G**), and the BC distributions in contact nasal turbinates and lungs became more similar over time (**Fig. S3H**). This provides additional evidence that SARS-CoV-2 can directly seed both the upper and lower airways in airborne-exposed hamsters and suggests that SARS-CoV-2 can readily disseminate throughout the respiratory tract.

### Hamster Sex does not Affect the Transmission Bottleneck

In the experiments described so far, all donor and contact hamsters were male. Male hamsters experience more weight loss and exacerbated lung injury compared to female hamsters after SARS-CoV-2 infection, though viral titers are similar between the sexes^49^. Similarly, male humans generally have greater disease severity than females^50^. To assess whether hamster sex affects the SARS-CoV-2 transmission bottleneck, we inoculated male and female donor hamsters with 10^5^ PFU of the BC pool and exposed sex-matched contact hamsters 24 hrs later in our airborne transmission model (**Fig. S4A**). Nasal turbinates, trachea, and lungs were collected from donor hamsters immediately after exposure and from contact hamsters 72 hpe, virus titers were measured by RT-qPCR, and BC distributions were measured by NGS. All tissues in all male and female contacts were positive for viral RNA by RT-qPCR, and hamster sex did not affect viral titers in donors (**Fig. S4B**) or contacts (**Fig. S4C**). Further, hamster sex did not affect the number of unique BCs transmitted to contacts, whether looking at whole hamsters or individual respiratory tissues (**Fig. S4D**). Additionally, the similarity between the nasal turbinates and lungs of individual hamsters was indistinguishable between males and females (**Fig. S4E**), and both sexes have BCs shared across all respiratory tissues and tissue-specific BCs (**Fig. S4F-G**). These results suggest that hamster sex has minimal impact on SARS-CoV-2 transmission. As such, male hamsters were used as donors and contacts in all subsequent experiments.

### Prolonged Exposure to Infected Donors Increases the Transmission Bottleneck

Duration of exposure to a SARS-CoV-2 infected individual has been associated with higher infection rates in both humans^51^ and animal models^52^. To measure how time of exposure impacts the transmission bottleneck in an airborne exposure setting, we inoculated donor hamsters with 10^5^ PFU of the BC pool and exposed contact hamsters 24 hrs later for 1, 4, or 8 hrs in our airborne transmission model (**Fig. 3A**). Nasal turbinates, trachea, and lungs were collected from contact hamsters 72 hpe, virus titer was measured by RT-qPCR, and BC were detected by NGS. Following a 1 hr exposure to SARS-CoV-2 infected donor hamsters, 50% (4/8) of the airborne-exposed contact hamsters were positive for SARS-CoV-2, while 100% of the hamsters exposed for 4 or 8 hrs were infected (**Fig. 3B**). Peak viral titers were generally similar across all groups (**Fig. 3C**). The average number of unique BC detected were 1, 7, and 15 following 1, 4 and 8 hour exposures, respectively, and all increases in exposure duration were associated with a statistically significant increase in BC counts (**Fig. 3D**). These results suggest that longer exposures not only increase the probability of infection, but also the number of viruses transmitted to the contact individual.

**Figure 3:**
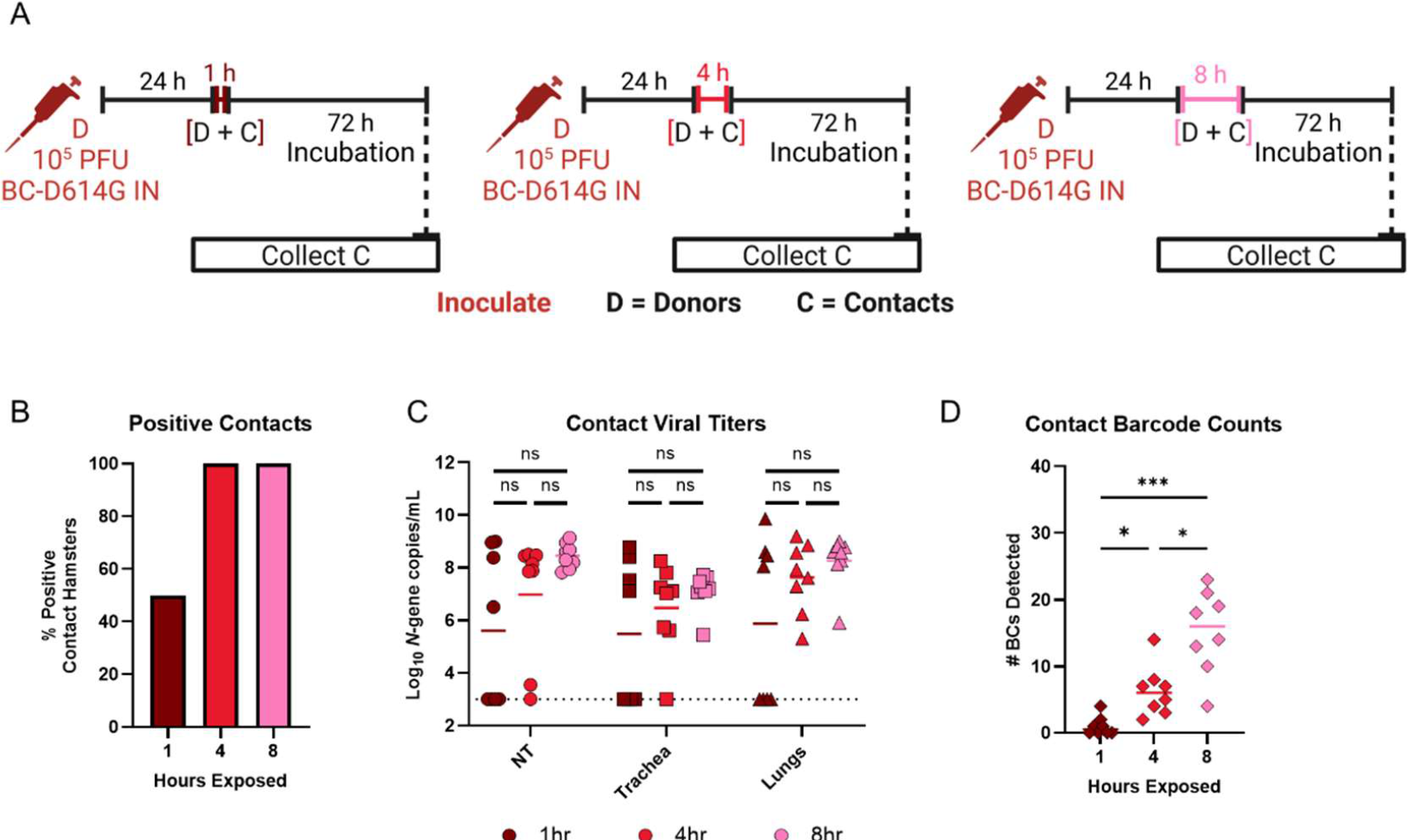
Prolonged exposure to SARS-CoV-2 increases transmission bottleneck in airborne-exposed hamsters. **(A)** Schematic of experimental design. Contact hamsters (n=8 per group) were airborne exposed for 1 (left, dark red), 4 (middle, red) or 8 (right, pink) hours to donor hamsters that were inoculated with 10^5^ PFU of the BC virus pool. Seventy-two hours later, nasal turbinate, trachea, and lung tissues were collected, and RNA extracted. (**B**) Fraction of airborne-exposed contact hamsters that were positive (RNA above the limit of detection) for SARS-CoV-2 by RT-qPCR in one or more tissues. (**C**) Virus titers, measured by RT-qPCR, in the nasal turbinate (circles), trachea (squares), and lung (triangles) of airborne exposed contact hamsters. Each symbol is an individual animal, and the line is the geometric mean of the data. Log-transformed data was analyzed by a Kruskal-Wallis test and a Dunn’s multiple comparison’s test. (**D**) The number of unique barcodes detected in contact hamsters (one-way ANOVA with multiple comparisons correction). Dotted lines indicated limit of detection. The results are from two independently repeated experiments with four donor and contact animals each. (*** *P* < 0.001, ** *P* < 0.01, * *P* < 0.05, ns = not significant).

### Route of Exposure Defines the Transmission Bottleneck

The impact of route on the transmission bottleneck is not well defined for many respiratory viruses including SARS-CoV-2. Here, we assessed the impact of route of exposure on the transmission bottleneck using the BC pool. Donor hamsters were inoculated IN with 10^5^ PFU of the BC pool. Twenty-four hours later, naïve contact hamsters were exposed to donor hamsters for 8hrs using the airborne or direct contact route. To model direct-contact transmission, donor and contact hamsters were placed in a clean cage without the 2cm barrier used for airborne-exposures. To model fomite transmission, naïve contact hamsters were placed in the dirty cage and bedding that had housed a previously infected donor animal for 8hrs before returning the contact to its original cage. Nasal turbinates, trachea, and lung were collected from contact hamsters 72 hpe (**Fig. 4A**), viral titers were measured by RT-qPCR, and the BC distribution in each tissue was measured by NGS. All contact animals, regardless of the route of exposure, were positive for SARS-CoV-2 (**Fig. 4B**) and viral titers within each tissue tested were similar between all three routes of exposure (**Fig. 4C**). However, there were substantial differences in the number of BCs detected across different routes of exposure. Fomite-exposed animals were infected with an average of 8 unique BCs. This number increased significantly (*P* < 0.05) to 16 unique BCs in airborne-exposed animals. Direct contact-exposed hamsters had the highest number of unique BCs detected (average 27). This frequency was significantly higher compared to fomite and airborne-exposed hamsters (**Fig. 4D**). Of note, tissue-specific BCs were identified in the upper and lower respiratory tract of animals from all routes of exposure (**Fig. 4E-F**). Collectively, these data suggest that more direct routes of exposure increase the SARS-CoV-2 transmission bottleneck.

**Figure 4:**
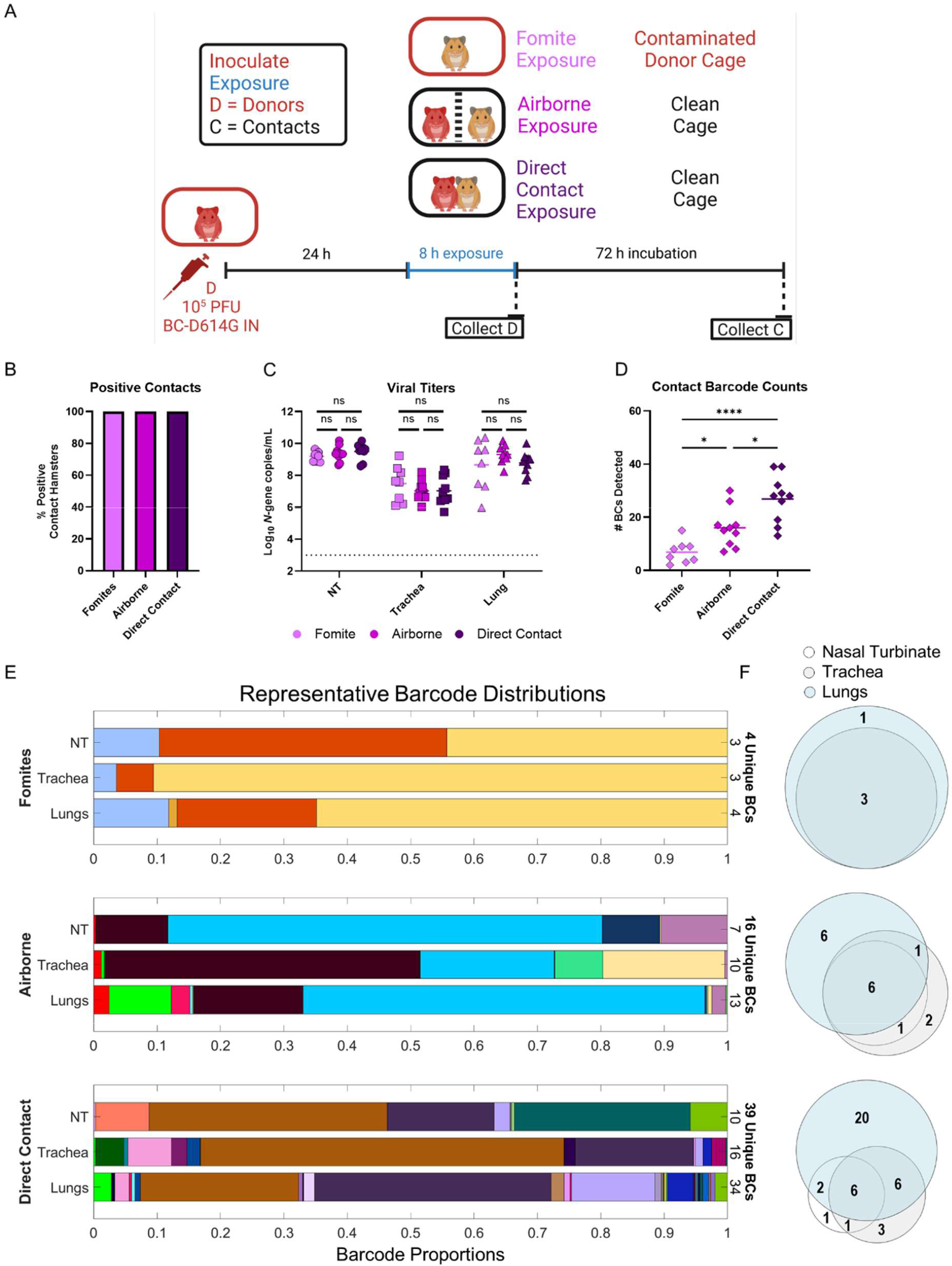
Route of exposure affects transmission bottleneck of SARS-CoV-2 in Syrian hamsters. **(A)** Schematic of experimental design. Contact hamsters (n=8-10 per group) were exposed to donor hamsters 24hrs after inoculation with 10^5^ PFU of the BC virus pool via fomite, airborne, or direct-contact route. Seventy-two hours later, nasal turbinate, trachea, and lung tissues were collected, and RNA extracted. (**B**) Fraction of SARS-CoV-2-positive contact hamsters. Transmission was defined as one or more tissues with viral RNA levels above the limit of detection. (**C**) Viral titers were measured by RT-qPCR in RNA extracted from nasal turbinate (circles), trachea (squares), and lung (triangles) of exposed contact hamsters. Dotted lines indicated limit of detection. Each symbol is an individual animal, and the line represents the geometric mean of the data. Data was analyzed by one-way ANOVA with multiple comparisons correction on log-transformed data. (**D**) The total number of unique barcodes detected across all three respiratory tissues in the fomite, airborne and direct-contact exposed hamsters. Each symbol is one animal and the bar represent the average transmission bottleneck (one-way ANOVA with multiple comparisons correction). (**E**) Representative barcode distributions for each route of exposure. Each color represents a unique BC and bar width is proportional to the BC’s relative frequency. The total number of unique BCs detected in each tissue and in the entire animal is shown on the right side of the graph. (**F**) Proportional Venn diagram showing barcodes unique to and shared between tissues in each animal. The results are from two independently repeated experiments. (**** *P* < 0.0001, *** *P* < 0.001, ** *P* < 0.01, * *P* < 0.05, ns = not significant).

## DISCUSSION

The SARS-CoV-2 transmission bottleneck is important to understand given its impact on SARS-CoV-2 evolution. In this study, we used a reverse genetics system to generate a pool of >200 BC SARS-CoV-2 viruses. Following in-depth characterization and validation of our BC SARS-CoV-2 pool, we used this pool to study SARS-CoV-2 transmission in hamsters, an important preclinical animal model. To this end, we infected donor hamsters with the entire pool and exposed contacts, varying the route and duration of exposure and measuring the impact on the number of BCs transmitted. Longer exposures in our airborne model widened the transmission bottleneck. We also defined the transmission bottleneck in other exposure models, with direct contact exposure having the widest bottleneck (average 27 BCs), followed by airborne exposure (average 16 BCs), followed by fomite exposure (average 8 BCs). Unique infection events were detected in the upper and lower airways of exposed hamsters, independent of time and route of exposure. Overall, these data demonstrate the power of BC viruses for understanding transmission and transmission bottlenecks of SARS-CoV-2.

A major technical advance of this work is the generation of a BC SARS-CoV-2 pool. Despite the impact of viruses in the family *Coronaviridae* on human and animal health, this is the first report to our knowledge of a recombinant barcoded coronavirus. Further, we demonstrated the quality our BC virus pool to a novel level of quantitative rigor, establishing a pipeline that future studies using BC viruses can follow.

While prior studies in hamsters have shown that fomite transmission is less efficient than airborne or direct contact^53^, potentially driven by the short half-life of SARS-CoV-2 on many materials^54^, this is the first study to our knowledge to quantitatively assess the transmission bottleneck in fomite, airborne, and direct contact settings, and to measure the impact of time of exposure on the transmission bottleneck. These results bolster public health guidance given throughout the pandemic, suggesting that not only do shorter and less direct exposures decrease the probability of infection, but also that they decrease the transmission bottleneck and thereby potentially the rate of SARS-CoV-2 evolution. Additionally, the finding that longer exposure times increases SARS-CoV-2’s transmission bottleneck suggests that SARS-CoV-2 can readily superinfect. Superinfections with different strains could enable recombination events, and several lines of evidence suggest that SARS-CoV-2 could leverage recombination to expedite its evolution. Many recombinant coronaviruses have been detected in nature^55^, mixed-strain SARS-CoV-2 infections have been observed in humans^56, 57^, and recombinant strains of SARS-CoV-2 have been reported in humans (e.g. Deltacron^58^, a recombinant between the Delta (B.1.617) and Omicron (BA.1) variants of SARS-CoV-2. Altogether, these reports highlight that SARS-CoV-2 may be capable of facile superinfection in humans, and that such superinfections could generate fit recombinant strains of SARS-CoV-2.

Following a 1 hr airborne exposure half of the contact animals became positive for SARS-CoV-2 by RT-qPCR (**Fig. 3B**). Interestingly, two of these animals were infected with a single BC that had disseminated across multiple respiratory tissues. This suggests that a single virion is sufficient to establish a productive infection in hamsters, a finding that is also supported in recent work using the Alpha and Delta variants^59^. Additionally, we saw no difference in titers across different routes of exposure, despite an almost 4-fold change in the transmission bottleneck. This implies that inoculating dose may have little impact on peak viral titers and potentially disease severity, a finding also supported in humans^60^.

Another advantage of BC viruses is the capacity to detect unique transmission events to different parts of the respiratory tract and follow subsequent dissemination. We detected unique BCs in the upper and lower airways of contacts 24, 48, and 72 hpe, with more tissue-specific BCs at earlier timepoints (**Fig S3G-H**). This strongly suggests that transmitted SARS-CoV-2 can not only seed the upper airways, but also bypass to directly seed the lungs, potentially transported by smaller respiratory particles. Further, the increase in shared BCs over time suggests efficient dissemination of SARS-CoV-2 throughout the respiratory tract. Future studies should investigate whether and how direct seeding of the lungs and efficient dissemination throughout the respiratory tract occur in humans, as these could have important implications for virus evolution.

These findings also highlight differences between SARS-CoV-2 and other respiratory viruses. As mentioned above, in influenza transmission, expansion in contacts is a stringent bottleneck^22^. We detected no evidence of a bottleneck during expansion in SARS-CoV-2 (**Fig S3**). Additionally, data using BC influenza suggests that influenza does not readily disseminate between the upper and lower respiratory tract^21, 23^. In contrast, as mentioned above, SARS-CoV-2 can readily disseminate between the upper and lower respiratory tract of hamsters. These results suggest that any intra-host SARS-CoV-2 variant may have a much greater capacity to disseminate from its site of infection to a site from which it could be transmitted, increasing the likelihood that it spreads to a new host.

Finally, these results have important implications for how to most accurately model human-to-human transmission using hamsters. As mentioned above, the transmission bottleneck in people has been estimated to be 1-5 viruses. The results shown here suggest some ways in which tracking intrahost variants may underestimate the transmission bottleneck. First, if SARS-CoV-2 can directly seed the lung in humans, there may be lung-specific variants that would be undetectable in nasal swabs. Second, intrahost variants must be present above a frequency cutoff (typically 1-3%) to be detectable, and we regularly detected BCs below this cutoff, suggesting that rare variants may be missed. However, we do not think these parameters will significantly impact estimations of the SARS-CoV-2 transmission bottleneck in people. For example, Bendall et al^17^ identify 64 transmission pairs, and only 6 transmission pairs (all from the same household) were shared multiple variants between the index case and contacts. As such, even if a majority of intrahost variants are undetected due to being present at a low frequency or lung-specific, the estimated transmission bottleneck for SARS-CoV-2 would remain narrow. Given the wide transmission bottlenecks measured here following 8 hr exposures, we would argue that either short (1-2 hour) airborne exposures or exposure models with greater physical distance separating hamsters (as in Port et al^59^) are likely more accurate models of human-to-human transmission than prolonged, direct exposure models.

The use of BC SARS-CoV-2 does come with several caveats. First, no BC viruses were whole-genome sequenced, so some BCs may harbor mutations elsewhere in the genome. However, we were unable to detect any fitness-enhancing mutations *in vitro* or *in vivo* and believe that rare fitness-detracting mutations in some BCs would not meaningfully impact the main conclusions of this paper. Second, the BC count can accurately reflect transmission bottleneck if all transmission events are independent; however, if SARS-CoV-2 is transmitted in particles containing multiple functional virions, as has been shown for rotavirus and norovirus^61^, the BC strategy will underestimate the transmission bottleneck, as it cannot distinguish between the transmission of a single BC virus and a multiple virions all harboring the same BC. The detection of single BCs in 50% of the 1 hr airborne-exposed contact hamsters, suggests that this is not the case in this model. Further, a recent modeling study^62^ suggests that it is unlikely that multiple respiratory viruses are transmitted in the same particle, but the question is not fully settled. Finally, the diversity of this BC pool is such that we can only detect 20-30 transmission events before there is an appreciable likelihood that multiple BCs are transmitted in duplicate. As such, the number of BCs detected in our 8hr airborne-exposed and direct contact-exposed hamsters represent lower limits of the true transmission bottleneck. Even with these caveats in view, BC- SARS-CoV-2 represents a significant technical advance which we have leveraged to study hamster-to-hamster SARS-CoV-2 transmission with unprecedented quantitative depth. BC- SARS-CoV-2 enabled experiments that defined of the transmission bottleneck under a range of routes and durations of exposure, identified direct infection events to the lung, which has implications for human transmission and disease, found evidence of facile superinfection, and elucidated important differences in virus transmission between SARS-CoV-2 and other respiratory viruses. This approach will add a novel and important dimension to future studies about how immunity impacts virus transmission.

## AUTHOR CONTRIBUTIONS

R.T. and T.L.D. performed all the *in vitro* and *in vivo* experiments. R.T. performed all the next-generation sequencing and barcode analysis. T.G. and D.W. designed the barcoded virus system. T.G., and T.L.D., and R.T. generated and expanded recombinant SARS-CoV-2 viruses containing unique genetic barcodes. A.C.M.B. had unrestricted access to all the data, analyzed the data, and performed the statistical analysis. D.W. and A.C.M.B. supervised experiments and acquired funding. R.T., T.L.D., and A.C.M.B. wrote the first draft of the manuscript and all authors reviewed and edited the final version. All authors agreed to submit the manuscript, read, and approved the final draft, and take full responsibility for its content.

## ACKNOWLEDGEMENTS

This study was supported by the NIH (NIAID Center of Excellence for Influenza Research and Response (CEIRR)) contract 75N93021C00016 (A.C.M.B.), R01AI169022 (A.C.M.B.) and U01AI151810 (A.C.M.B and D.W.).

## METHODS

***Cells.*** Vero cells expressing human angiotensin converting enzyme 2 (ACE2) and transmembrane serine protease 2 (TMPRSS2) (Vero-hACE2-hTMPRSS2, gift of Adrian Creanga and Barney Graham, NIH) were cultured at 37°C in Dulbecco’s Modified Eagle medium (DMEM) supplemented with 10% fetal bovine serum (FBS), 10 mM HEPES (pH 7.3), 2 mM L-glutamine, 100 U/mL Penicillin, 100 µg/mL Streptomycin, and 10 µg/mL of puromycin. Vero E6 cells expressing human TMPRSS2 (Vero-hTRMPSS2) were cultured at 37°C in DMEM supplemented with 10% FBS, 10 mM HEPES (pH 7.3), 2 mM L-glutamine, 100 U/mL Penicillin, 100 µg/mL Streptomycin, and 5 µg/mL of blasticidin. BHK-21 cells were cultured at 37°C in DMEM supplemented with L-glutamine, 10% FBS, 100 U/mL Penicillin, and 100 µg/mL Streptomycin. Vero-hACE2-hTRMPSS2 are used to titrate stocks and tissues, Vero-hTRMPSS2 cells are used to generate virus stocks and BHK-21 cells were used to generate barcoded virus.

### Recombinant barcoded SARS-CoV-2

We used our SARS-CoV-2 reverse genetics system, described previously^1^, to generate barcoded SARS-CoV-2 viruses containing the spike D614G mutation (**Fig. 1A**). Briefly, the SARS-CoV-2 genome (GenBank accession no. NC_045512) was split into 7 fragments (A-G), commercially synthesized as DNA (GenScript), and cloned into plasmids. A T7 promoter was added to the 5’ end of the viral fragment in plasmid A. We introduced the D614G mutation in the S gene in fragment F using the primers listed in Table 1. We inserted a 9nt barcode sequence of GNNGNNGNN through a degenerate primer (Table 1) into ORF10 cassette (after 21 nucleotide) in fragment G. Transformed competent cells were spread on LB plates and colonies scraped and pooled for propagation and plasmid extraction. The 7 plasmids, corresponding to fragments A-G, were digested with type II restriction enzymes (NEB), and the viral genome fragments were ligated with T4 ligase (NEB). The purified ligation product was used as the template for *in vitro* transcription to produce full-length viral genome using mMESSAGE mMACHINE T7 Ultra Kit (Invitrogen).

**Table 1.**
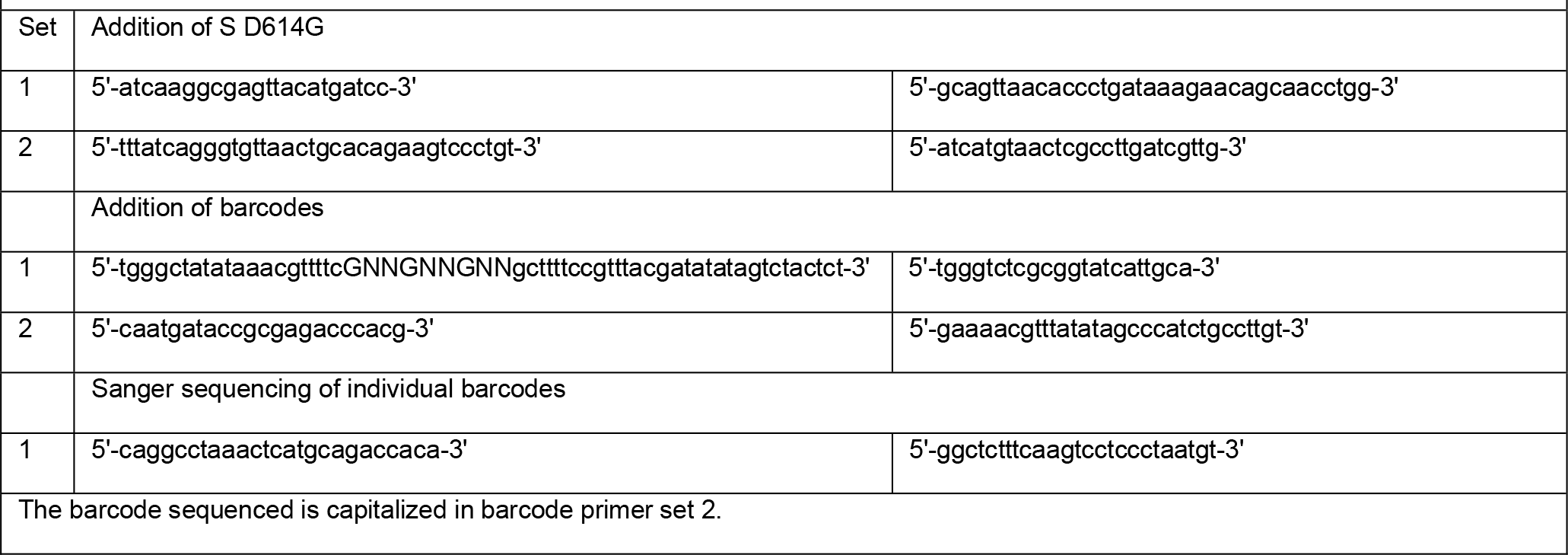
Primer sets for addition of S D614G and barcodes

The *in vitro*-transcribed barcoded viral genomes were electroporated into BHK-21 cells together with *N* gene mRNA following an established protocol^39^. Electroporated BHK-21 cells were diluted and mixed 1:1 with Vero-hACE2-hTMPRSS2 and distributed over several 24-well plates to 10^3.4^ cells per well. Cells were monitored daily for cytopathic effects (CPE) and supernatant was harvested from positive CPE wells 5 days post electroporation. Positive wells were expanded on Vero-hTMPRSS2 cells for 24hrs, aliquoted, and stored at -80°C. The infectious titers were measured by plaque assay on Vero-hACE2-hTMPRSS2 cells. To identify the barcode (BC) sequence, viral RNA was extracted from supernatant using an E.Z.N.A. Total RNA Kit (Omega), and used to amplify a 204bp amplicon in a one-step RT-PCR reaction (SuperScript IV One-Step, Thermo Fisher) using custom primers (Table 1). This amplicon was then Sanger sequenced using the reverse primer. All expanded samples that contained unique BC sequences were pooled based on the amount of infectious virus, aliquoted, and stored at -80°C for the studies in this manuscript. The relative frequency of each BC in pool was determined by next-generation sequencing and analysis as described below.

All work with infectious SARS-CoV-2 was performed in Institutional Biosafety Committee approved BSL-3 and ABSL3 facilities at Washington University School of Medicine using appropriate positive pressure air respirators and protective equipment.

*Hamster experiments*. Animal studies were carried out in accordance with the recommendations in the Guide for the Care and Use of Laboratory Animals of the National Institutes of Health. The protocols were approved by the Institutional Animal Care and Use Committee at the Washington University School of Medicine (assurance number A3381–01). Five-six week old male hamsters were obtained from Charles River Laboratories and housed at Washington University.

*Direct challenge*. Five-to-six week old male or female donor hamsters were inoculated with 10^5^ PFU of the BC virus pool. Thirty-two or seventy-two hrs later, hamsters were euthanized, and respiratory tissues were collected for virological and BC frequency analysis.

*Primary transmission.* Five-to-six week old male donor hamsters were inoculated with 10^5^ PFU of the BC virus pool and, 24hrs later, contact hamsters were exposed to donor hamsters for 1, 4 or 8hrs. For direct contact exposure, both donor and contact hamsters were directly placed in a clean biocontainment unit (BCU) cage with fresh bedding. For airborne exposure, donor and contact hamsters were placed in separate porous stainless steel (isolator) cages which were then placed in a single clean BCU cage with directional airflow from the donor to the contact hamster. For fomite exposure, contact hamsters were placed in the contaminated BCU cage from donor hamsters. After the exposure the donor and contact hamster were placed back into their original cage. Contact hamsters were euthanized, and tissues collected 24, 48, or 72hrs after exposure. Exposure and tissue collection times are indicated in the results section for each experiment.

At time of tissue collection, nasal turbinate, trachea, whole lung, or individual lung lobes were homogenized in 1.0 mL of DMEM and clarified by centrifugation at 1,000 x g for five minutes. To collect nasal turbinate, the skin along the nose and cheeks and the lower jaw were removed to expose the upper palate. A sagittal incision through the palate exposed the nasal turbinates, which were removed using blunt forceps. RNA was extracted and viral titer was determined by quantitative RT-qPCR and RNA sequenced in the barcode region to detect unique barcodes.

### Virus titration assays

Plaque assays were performed on Vero-hACE2-hTRMPSS2 cells in 24-well plates. Samples were serially diluted in cell infection medium (DMEM supplemented with 2% FBS, 10 mM HEPES (pH 7.3) and 2 mM L-glutamine). Two hundred microliters of the diluted virus were added to a single well per dilution per sample. After one hour at 37°C, the inoculum was aspirated, the cells were washed with PBS, and a 1% methylcellulose overlay in MEM supplemented with 2% FBS was added. Seventy-two hours after virus inoculation, the cells were fixed with 10% formalin, and the monolayer was stained with crystal violet (0.5% w/v in 25% methanol in water) for one hour at 20°C. The number of plaques were counted and used to calculate the plaque forming units/mL (PFU/mL). Infectious virus titer detected in any of the contact hamster organs was considered a positive transmission event.

To quantify viral RNA load in respiratory organ tissue homogenates, RNA was extracted using a modified protocol for the MagMax Viral Pathogen Kit on the KingFisher Flex (Thermo Fisher). Briefly, 100 µL clarified homogenate sample was lysed in 300 µL TRK lysis buffer (Omega) plus 2% βME. Samples were Proteinase K treated and then total nucleic acid magnetic beads were added in binding solution. The magnetic beads were then washed in wash buffer provided by the kit and twice in 80% Ethanol. Finally, RNA was eluted with 50 µL of water. Four microliters RNA was used for real-time RT-qPCR to detect and quantify genomic RNA of SARS-CoV-2 *N* gene using TaqMan™ RNA-to-CT 1-Step Kit (Thermo Fisher Scientific) as described^2^ using the following primers and probes: Forward: GACCCCAAAATCAGCGAAAT; Reverse: TCTGGTTACTGCCAGTTGAATCTG; Probe: ACCCCGCATTACGTTTGGTGGACC; 5’Dye/3’Quencher: 6-FAM/ZEN/IBFQ. Units are described in genome equivalent copy numbers per µL of RNA based on a standard included in the assay, which was created via *in vitro* transcription of a synthetic DNA molecule containing the target region of the *N* gene.

### Barcode analysis

*Sequencing.* To sequence the barcode distribution in samples with detectable SARS-CoV-2 genomic RNA by RT-qPCR, extracted RNA was used as a template in a cycle-limited one-step RT-PCR reaction (SuperScript IV One-Step, Thermo Fisher) using custom primers (Table 2, Set 1). The resulting amplicon was then used as a template in a PCR reaction (Q5, NEB) to add ends compatible with sequencing using custom primers (Table 2, Set 2). Libraries from different samples were pooled, purified using a QIAquick PCR purification kit (Qiagen), and sequenced (2x150 bp) on a Miniseq system (Illumina).

**Table 2.**
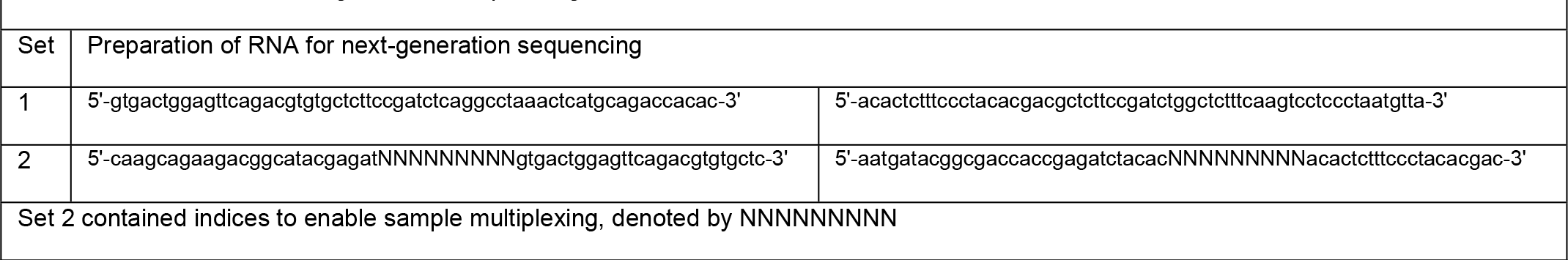
Primer sets for next-generation sequencing of barcode distributions

Sequencing data was demultiplexed and trimmed of adapter and index sequences. The barcode distributions in each sample were enumerated using custom code adapted from Weger-Lucarelli et al^34^. Briefly, paired-end reads were merged using BBMerge, aligned to the SARS-CoV-2 genome using BBMap, trimmed to the barcode region using Reformat.sh, and barcodes were counted using kmercountexact.sh. All of these programs are part of the BBTools suite^63^.

*Analysis of barcode distributions.* All analyses of barcode distribution data were performed using custom Matlab scripts, made publicly available at https://github.com/rtrende/BC_SARS-CoV-2_Analysis. Briefly, the barcode matrix was trimmed to just contain barcodes present in the inoculum. A barcode was deemed present in a sample if >0.056% (∼1/1800^th^) of the reads in that sample mapped to the barcode. This cutoff was set based on the frequency of mutants in known barcode pools (data not shown). Richness was calculated as the number of barcodes present above this threshold. Diversity (*H’*) was calculated using Equation 1, where *n* is the total number of BCs and *xi* is the relative frequency of the *i*-th BC:

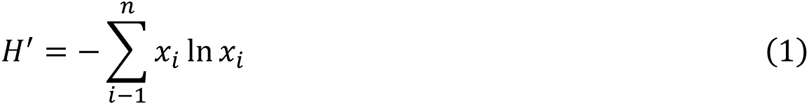

Evenness was calculated using Equation 2:

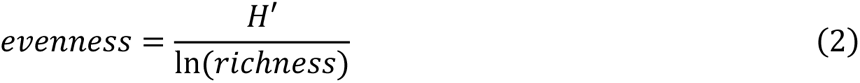

The similarity index (equivalent to “genetic relatedness” in ^41–43;^ based on Cavalli-Sforza chord distance^64^) was calculated using Equation 3, where *xi* and *yi* represent the relative frequency of the *i*-th BC in the two samples between which similarity is calculated:

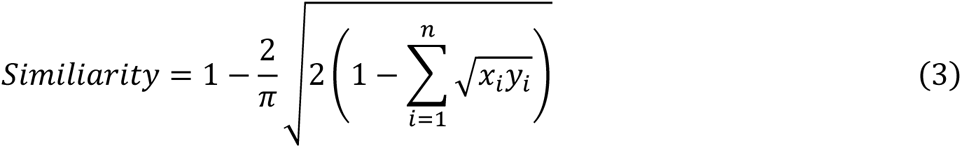

*Analysis of infectious BC virus pool from tissue homogenates.* Approximately 10^6^ PFU the BC virus pool was amplified from nasal turbinate homogenate on Vero-TMPRSS2 cells at an MOI = 0.01. After 2 h, inoculum was aspirated and cells washed three times with PBS. Three mL infection medium was added per well. Supernatant was collected at 24 and 48 hpi. RNA was extracted and the barcode region was sequenced.

### Quantification and statistical analysis

Statistical significance was assigned when P values were < 0.05 using GraphPad Prism version 10.1. Tests, number of animals, and statistical comparison groups are indicated in the Figure legends. When two groups were compared, data were analyzed using unpaired t-tests followed by a Holm-Šídák test; when more than two groups were compared, data were analyzed using a Brown-Forsythe and Welch ANOVA followed by Dunnett’s T3 multiple comparisons test. Viral titers were log-transformed prior to analysis.

---

**Figure S1:**
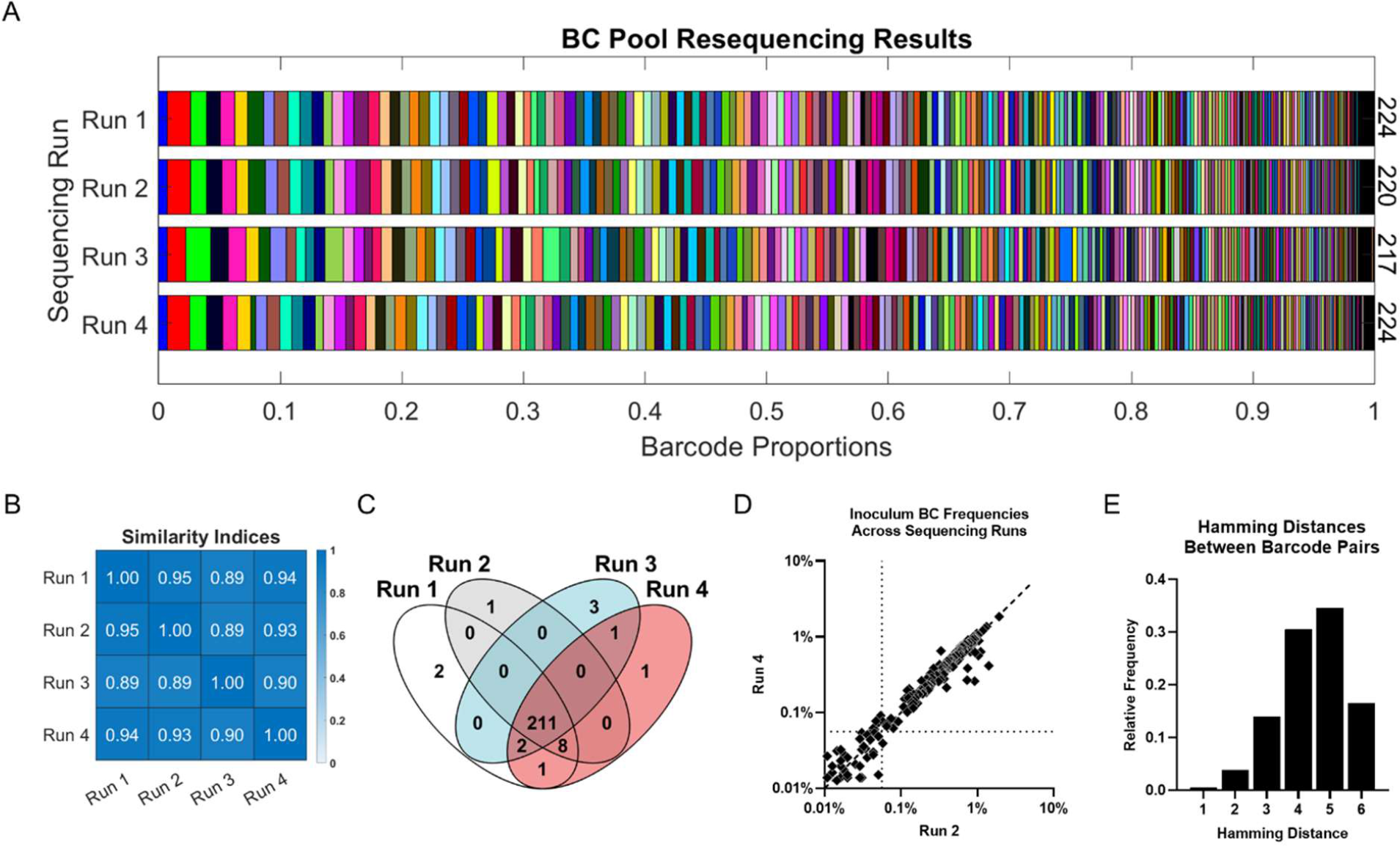
Pipeline for measuring distribution of BC pool yields highly reproducible results **(A)** RNA was extracted from 4 separate aliquots of the BC virus pool, and BC sequence and frequency was determined by NGS. Each color represents a unique BC and bar width is proportional to the BC’s relative frequency. The total number of unique BCs detected in each sample is shown on the right side of the graph. (**B**) Heatmap of similarity indices between each sequencing run. (**C**) Venn diagram showing overlap between the BCs detected in different sequencing runs of the BC pool. (**D**) Correlation of BC frequencies between two representative BC sequencing runs. (**E**) Distribution of Hamming Distances, i.e. the number of nucleotide difference, separating all BC pairs in the BC virus pool.

**Figure S2:**
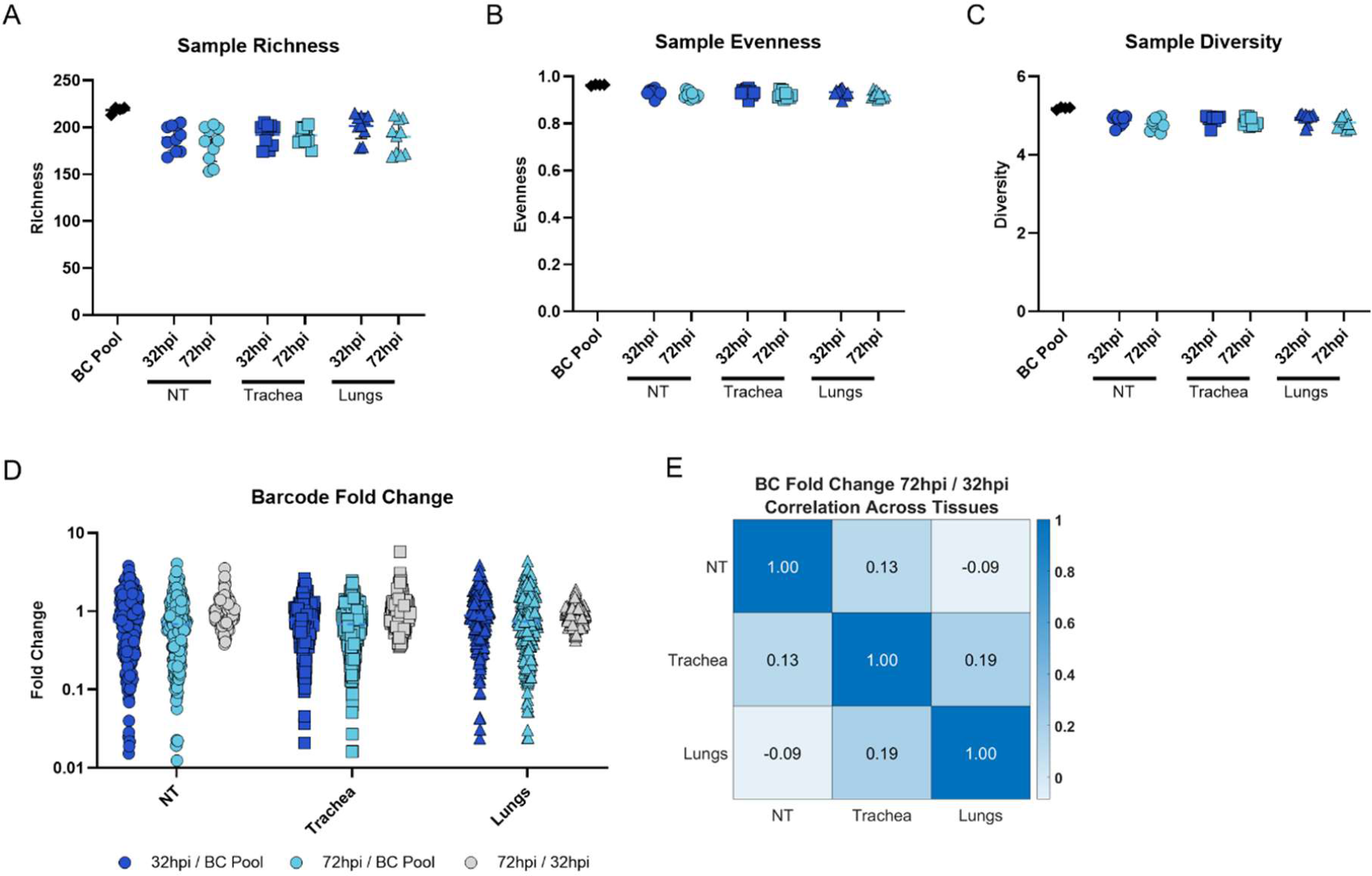
BC Pool Diversity is retained in vivo. **(A)** Richness, (**B**) evenness, and (**C**) Shannon diversity of BC distributions in inoculum and each hamster tissue collected 32 or 72 hours after intranasal inoculation with 10^5^ PFU of the BC virus pool. Each symbol is an individual hamster. The line represents the average of the data. (**D**) The geometric mean fold change in the relative BC frequency between the inoculum and the 32 or 72hpe timepoint, or between 32 and 72hpe time-point, in the nasal turbinate, trachea and lungs of inoculated hamsters. (**E**) Correlation matrix showing the Pearson correlation coefficient between geometric mean BC fold changes from 32hpi to 72hpi in the indicated tissues. Strong correlation indicates true differences in viral fitness, while weak correlation suggests that most changes in BC frequency are due to noise rather than fitness differentials. The results are from 2 independently repeated experiments with five hamsters each.

**Figure S3:**
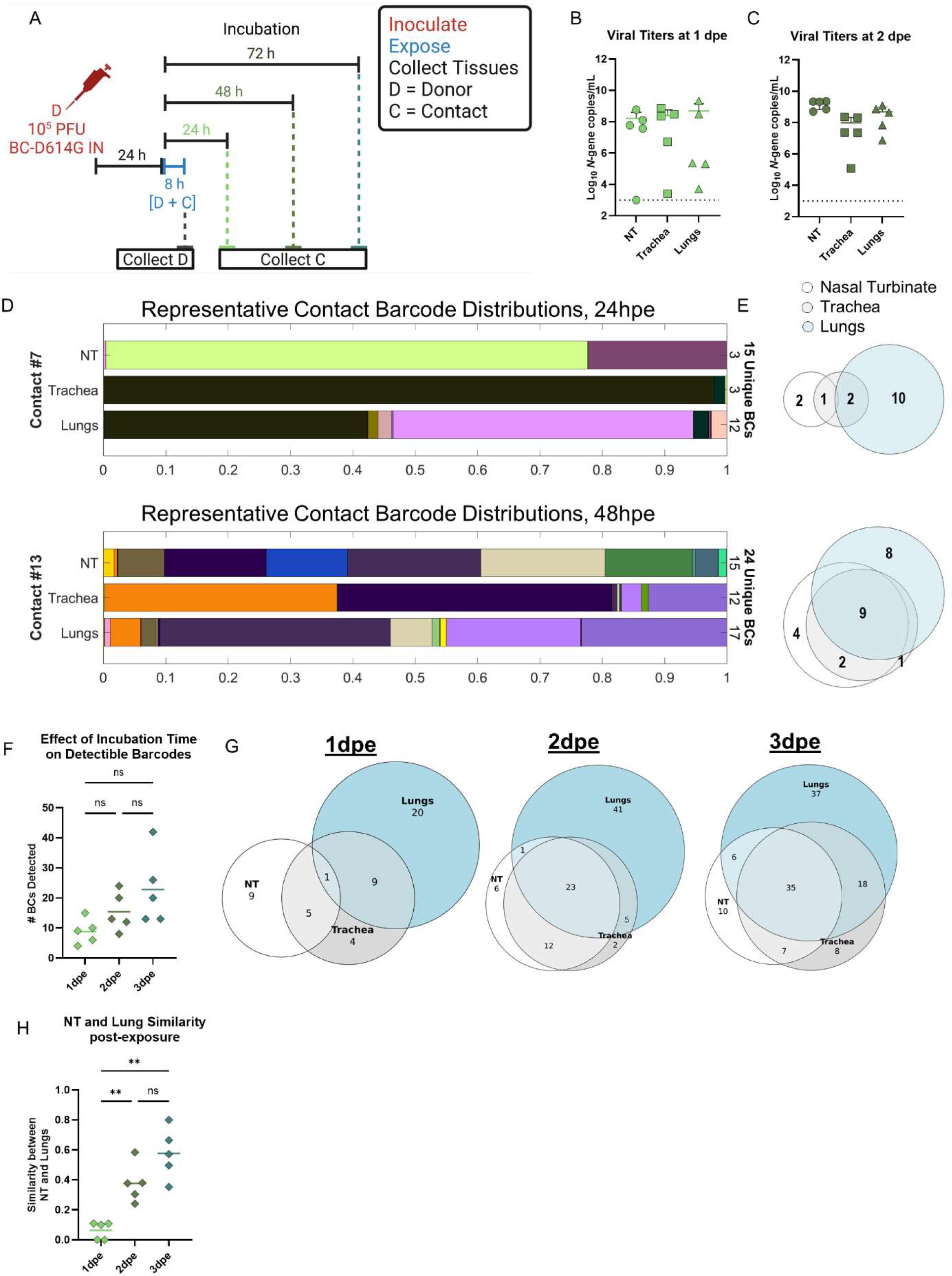
Defining the SARS-CoV-2 airborne transmission bottleneck 24-72 hours post exposure. **(A)** Schematic of experimental design. Contact hamsters (n=5) were airborne exposed to donor hamsters 24 h after inoculation with 10^5^ PFU of pooled barcoded SARS-CoV-2. Twenty-four, or forty-eight hours later, nasal turbinate, trachea, and lung tissues were collected, and RNA extracted. (**B** and **C**) Virus titers were measured by RT-qPCR, in nasal turbinate (circles), trachea (squares), and lungs (triangles) of airborne-exposed hamsters. Dotted lines indicated limit of detection. Each symbol is an individual animal and line indicates geometric mean. (**D**) Representative images BC distributions in contact hamsters collected 24 or 48hpe. Each color represents a unique BC and bar width is proportional to the BC’s relative frequency. The total number of unique BCs detected in each tissue and in the entire hamster is shown on the right side of the graph. (**E**) Proportional Venn diagram showing the number of unique barcodes in each tissue and shared between the tissues of representative hamsters 24 or 48hpe. (**F**) Total number of unique BC detected per hamster 24, 48, or 72hpe (see also Figure 2). (**G**) Proportional Venn diagram showing the number of unique and shared BCs in each tissue. Data shown is pooled from all hamsters at each timepoint. (**H**) Similarity indices between the nasal turbinates and lungs of contact hamsters collected at different timepoints. Each symbol is an individual hamster, and the line represents the average number of BC detected. Data are analyzed by Brown-Forsythe and Welch ANOVA followed by Dunnett’s T3 multiple comparisons test. The results are from a single experiment. (**** *P* < 0.0001, *** *P* < 0.001, ** *P* < 0.01, * *P* < 0.05, ns = not significant).

**Figure S4:**
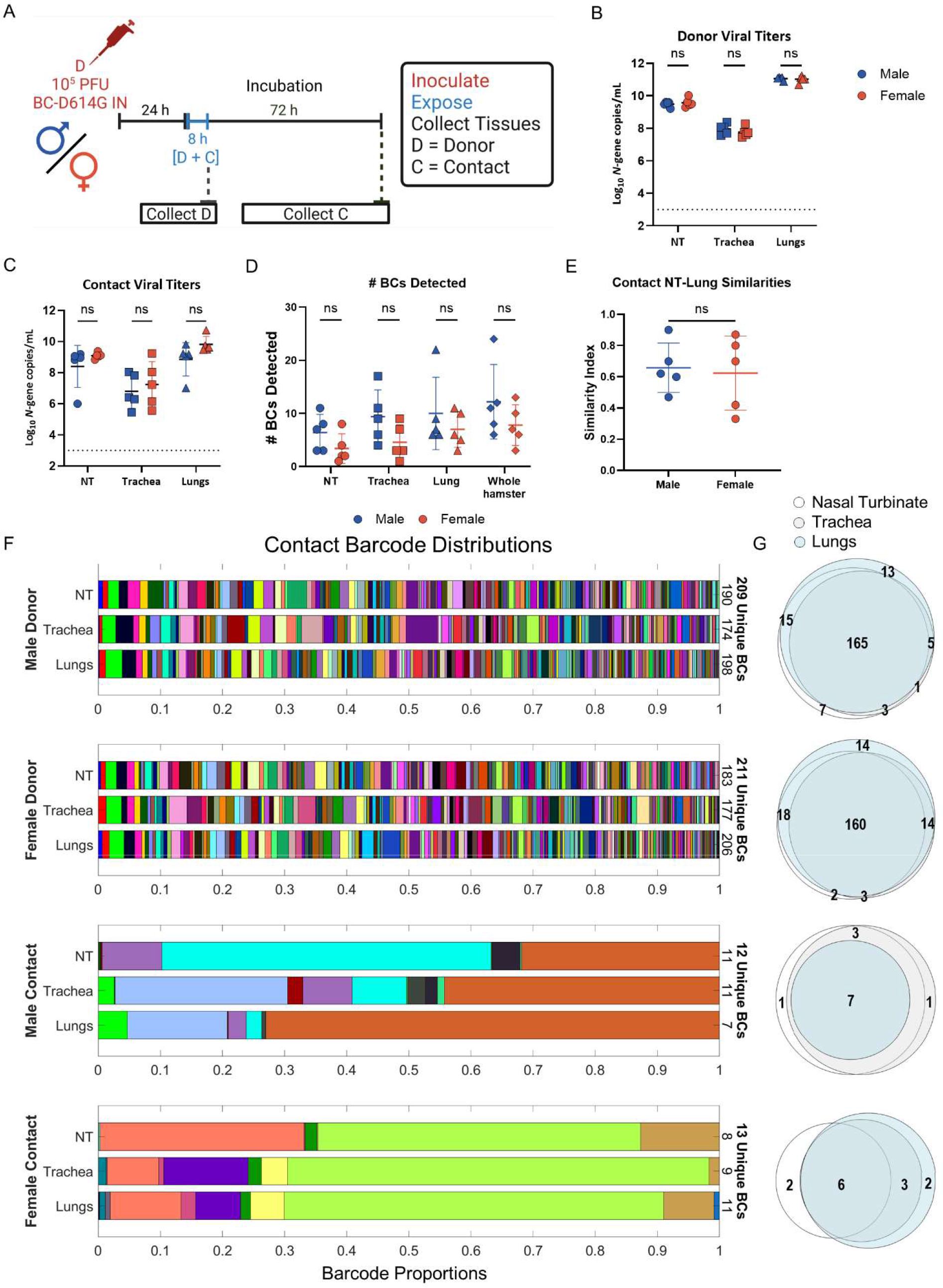
Hamster sex does not impact the SARS-CoV-2 transmission bottleneck. **(A)** Schematic of experimental design. Male and female donor hamsters (n=5 per group) were inoculated with 10^5^ PFU of the BC virus pool. Twenty-four hours after inoculation, sex-matched contact hamsters were airborne exposed for 8 hrs. Donor nasal turbinate, trachea, and lungs were collected immediately after exposure, while contact tissues were collected 3dpe. (**B**) Viral titers were measured by RT-qPCR in RNA extracted from the lungs of donor and contact hamsters. Dotted line indicates limit of detection. Each symbol is an individual animal, and the line represents the geometric mean. (**C**) The total number of unique BCs detected in individual tissues and in the entire animal from contact hamsters. Each symbol is one animal, and the bar represents the mean. Data was analyzed by unpaired t-tests. (**D**) Similarity index between the BC distributions in the nasal turbinate and the lungs of contact male and female hamsters. Symbols represent individual hamsters, lines represent mean. (**E**) Representative barcode distributions for each route of exposure. Each color represents a unique BC and bar width is proportional to the BC’s relative frequency. The total number of unique BCs detected in each tissue and in the entire animal is shown on the right side of the graph. (**F**) Proportional Venn diagram showing barcodes unique to and shared between tissues in each animal. Data was log-transformed when necessary and analyzed by unpaired t-tests followed by a Holm-Šídák test. (**** *P* < 0.0001, *** *P* < 0.001, ** *P* < 0.01, ns = not significant).

